# White Spot Syndrome Virus Immediate-Early Protein (wsv100) Antagonizes the NF-κB Pathway to Inhibit Innate Immune Response in shrimp

**DOI:** 10.1101/2024.12.16.628618

**Authors:** Bang Xiao, Fang Kang, Qianqian Li, Junming Pan, Yue Wang, Jianguo He, Chaozheng Li

## Abstract

Viruses have evolved sophisticated strategies to evade host immune defenses, often targeting conserved signaling pathways. In shrimp, the NF-κB signaling pathway is crucial for antiviral immunity, yet its regulation during White Spot Syndrome Virus (WSSV) infection remains poorly understood. Here, we identify and characterize wsv100, an immediate-early (IE) protein of WSSV, as a key antagonist of the NF-κB pathway. wsv100 interacts directly with the transcription factor Dorsal and the adaptor protein IMD, preventing Dorsal phosphorylation by Pelle kinase. This inhibition suppresses Dorsal’s nuclear translocation and downstream expression of antimicrobial peptides (AMPs), essential for antiviral defense. Knockdown of wsv100 reduced WSSV replication, increased Dorsal phosphorylation, and enhanced AMP expression, leading to higher survival rates in infected shrimp. Conversely, wsv100 overexpression promoted WSSV replication and AMPs suppression. These findings reveal a novel immune evasion mechanism by which WSSV subverts the NF-κB pathway and highlight the evolutionary arms race between hosts and viruses. This study enhances our understanding of host-virus interactions and offers potential targets for antiviral strategies in shrimp aquaculture.

**Author Summary:** The innate immune system is the first line of defense against viral infections in invertebrates, with the NF-κB signaling pathway playing a central role in orchestrating antiviral responses. In this study, we uncover a novel immune evasion mechanism employed by White Spot Syndrome Virus (WSSV), a devastating pathogen in shrimp aquaculture. The WSSV immediate-early protein wsv100 directly targets the transcription factor Dorsal and prevents its phosphorylation by Pelle kinase, a critical step in NF-κB activation. This interaction suppresses Dorsal’s nuclear translocation and downstream expression of antimicrobial peptides (AMPs), thereby impairing the shrimp’s ability to mount an effective immune response. Knockdown of wsv100 significantly reduced WSSV replication and enhanced shrimp survival, while wsv100 overexpression had the opposite effect. These findings not only elucidate how WSSV exploits the NF-κB pathway but also underscore its central role in shrimp antiviral immunity. This work advances our understanding of host-virus co-evolution and provides a foundation for developing novel antiviral strategies to mitigate the economic losses caused by WSSV in shrimp aquaculture.

## 1. Introduction

Viruses and their hosts are engaged in a perpetual evolutionary arms race, wherein hosts develop intricate immune defenses to counteract viral infections, while viruses evolve sophisticated mechanisms to evade or exploit these defenses [1, 2]. This dynamic interplay drives the co-evolution of host immune systems and viral counter-defense strategies, leading to increasingly refined molecular mechanisms on both sides [1]. In invertebrates, the innate immune system serves as the primary line of defense against pathogens, as these organisms lack adaptive immunity [3]. Central to this defense are evolutionarily conserved signaling pathways, including Toll, immune deficiency (IMD), JAK-STAT, and RNA interference (RNAi) [4–6]. Traditionally, the Toll and IMD pathways have been associated with antibacterial defenses: the Toll pathway primarily targets fungi and Gram-positive bacteria, while the IMD pathway targets Gram-negative bacteria [7]. However, emerging evidence suggests these pathways also play pivotal roles in antiviral immunity, challenging conventional paradigms and opening new avenues for research into host-virus interactions.

Among the signaling pathways implicated in host-virus interactions, the NF-κB signaling network in invertebrates, encompassing the Toll and IMD pathways, has emerged as a compelling system for study [8]. Both pathways play central roles in detecting pathogens and activating downstream immune responses. The canonical Toll pathway involves the extracellular ligand Spätzle (Spz), Toll receptors, adaptor proteins Tube and MyD88, kinases such as Pelle, and transcription factors Dorsal and Dorsal-related immunity factor (Dif) [9–11]. Similarly, the IMD pathway includes pattern recognition receptors (PGRPs), the adaptor IMD, kinases such as TAK1 and IKKβ, and the transcription factor Relish [12–16]. Upon viral infection, both pathways activate signaling cascades that lead to the nuclear translocation of Dorsal and Relish, triggering the production of antimicrobial peptides (AMPs) that combat infection [17]. This convergence of pathways in AMPs production underscores their central role in innate immunity and highlights their potential as critical antiviral defenses.

Experimental evidence increasingly supports the antiviral functions of the Toll and IMD pathways in invertebrates. In *Drosophila*, for example, deletion or silencing of Toll pathway components such as Toll, Spätzle, Pelle or Dorsal significantly heightens susceptibility to viral infections, including *Drosophila C virus* (DCV) [18]. Similarly, disruption of the IMD pathway by targeting key molecules like IKKβ or Relish impairs the host’s ability to control viral replication [19]. In shrimp, RNAi-mediated knockdown of Toll pathway components such as Toll4 results in increased replication of White spot syndrome virus (WSSV) and elevated host mortality [20]. These findings illustrate the crucial antiviral roles of the Toll and IMD pathways and emphasize the evolutionary pressures viruses exert to subvert these defenses.

Significantly, a key criterion for determining the antiviral function of these pathways is the identification of virus-encoded proteins that specifically block or hijack them [21]. Such viral strategies provide direct evidence of the pathways’ importance in host defense and reveal the molecular arms race between viruses and their hosts. In vertebrates, the antiviral role of NF-κB pathways is well established. Viruses such as Kaposi’s sarcoma-associated herpesvirus (KSHV), herpes simplex virus (HSV) and severe acute respiratory syndrome coronavirus 2 (SARS-CoV-2) have evolved mechanisms to inhibit NF-κB signaling, underscoring its critical role in host defense. For instance, KSHV’s transcription activator protein (RTA) disrupts the localization of Toll-like receptors (TLR2/4), thereby inhibiting downstream signaling, while HSV’s US3 protein blocks the nuclear translocation of p65, a central transcription factor [22, 23]. Additionally, the NSP8 and glycoprotein M protein of SARS-CoV-2 significantly inhibit the phosphorylation of TBK1, IRF3 and p65, thereby downregulating innate immune responses [24, 25]. These examples provide a framework for investigating similar interactions in invertebrates, where the antiviral roles of NF-κB pathways remain underexplored.

Shrimp, as aquatic invertebrates, rely on their innate immune system to fend off a diverse array of pathogens, including viruses, bacteria, and fungi [26]. WSSV, the sole member of the genus *whispovirus*, is one of the most devastating pathogens in shrimp aquaculture, causing widespread mortality and severe economic losses globally [27, 28]. To establish infection, WSSV employs diverse immune evasion strategies, including targeting host immune pathways. Among its 21 identified immediate-early (IE) genes, which are key regulators expressed at the onset of infection, wsv100 has emerged as a particularly intriguing candidate [29, 30]. The wsv100 gene encodes a protein featuring a coiled-coil domain, a TAZ zinc-finger domain, and nuclear localization signals, suggesting a role in protein-protein interactions and subcellular trafficking. While previous studies have highlighted the involvement of IE genes in viral replication and immune modulation, the precise mechanisms by which wsv100 facilitates immune evasion remain unclear.

In this study, we demonstrate that wsv100 specifically targets the NF-κB pathway to evade its antiviral function. wsv100 binds to Dorsal, preventing its phosphorylation by Pelle and subsequent nuclear translocation, thereby impairing the shrimp’s antiviral response. Furthermore, wsv100 interacts with IMD, an adaptor molecule in the IMD pathway, underscoring the virus’s ability to manipulate multiple branches of the NF-κB signaling network. From a virological perspective, these findings provide additional evidence supporting the antiviral role of NF-κB pathways, particularly the Toll pathway, in invertebrates. These insights deepen our understanding of invertebrate immunity, illuminate the molecular arms race between viruses and their hosts, and offer potential directions for developing antiviral strategies in aquaculture.

## 2. Results

### 2.1 Expression Profile of wsv100 During WSSV Infection

WSSV immediate-early (IE) genes play a pivotal role in viral infection and replication, with wsv100 (GenBank accession No. AWQ63223) being one of the 21 identified IE genes. However, the mechanisms by which WSSV employs wsv100 to evade the shrimp innate immune response remain unclear. The wsv100 protein contains a predicted coiled-coil domain, a TAZ zinc-finger domain, and two nuclear localization signals (Supplementary Fig. 1). The expression pattern of wsv100 in hemocytes and gills of WSSV-infected shrimp was similar to the IE1 gene (Figs. 1A, 1B). Consistent with the transcriptional data, western blot analysis revealed increased wsv100 protein levels in these tissues during infection (Figs. 1C, 1D). Immunofluorescence assays showed that wsv100 was predominantly cytoplasmic in hemocytes at 12 hours post-infection (hpi) but translocated to the nucleus by 24 hpi (Fig. 1E). These findings demonstrate that wsv100 expression and nuclear translocation are significantly upregulated during WSSV infection.

**Figure 1.**
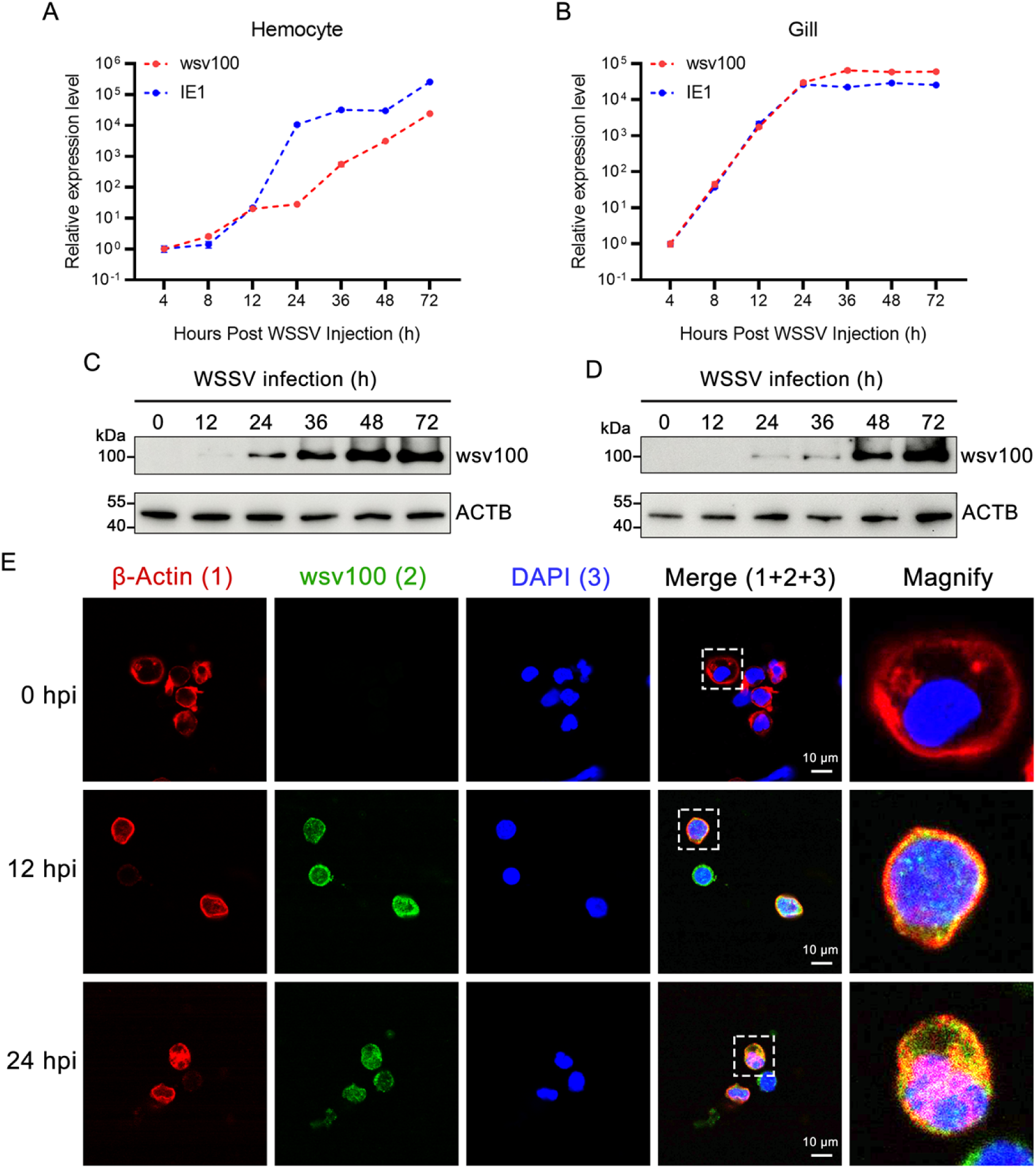
Expression profile of *wsv100* in WSSV-infected shrimp. (A-B) Quantitative PCR (qPCR) analysis of *wsv100* and *IE1* expression levels in hemocytes (A) and gills (B) following WSSV infection. (C-D) Western blot analysis of *wsv100* protein expression in hemocytes (C) and gills (D) after WSSV infection. (E) Immunofluorescence analysis showing the localization of *wsv100* (green) in hemocytes from WSSV-infected shrimp at 0, 12, and 24 hours post-infection (hpi). Actin (red) was used as a marker for the cell membrane, and Hoechst 33258 (blue) was used for nuclear staining. Scale bar = 10 μm. Statistical significance was determined using Student’s *t*-test (\*\**p* < 0.01; \**p* < 0.05). All experiments were performed in triplicate, and similar results were observed across biological replicates.

### 2.2 Knockdown of wsv100 Suppresses WSSV Replication

To explore the role of wsv100 during WSSV infection, RNA interference (RNAi) was used to knock down its expression *in vivo*. qRT-PCR confirmed effective silencing of wsv100 mRNA in hemocytes and gills at 24 and 48 hpi, respectively (Figs. 2A, 2B). Western blot analysis further showed reduced wsv100 protein levels at 48 hpi in both tissues (Figs. 2a, 2b). Notably, wsv100 silencing significantly lowered WSSV viral loads in hemocytes and gills at 24, 48, and 72 hpi compared with control shrimp injected with GFP-targeted dsRNA (Figs. 2C, 2D). The transcriptional and protein levels of VP28, a viral marker, were also reduced in wsv100-silenced shrimp (Figs. 2E, 2F, 2f). Furthermore, survival analysis revealed a significantly higher survival rate in wsv100-knockdown shrimp compared to the GFP-silenced controls (*p* < 0.001; Fig. 2G). These results strongly suggest that wsv100 knockdown suppresses WSSV replication and enhances host survival.

**Figure 2.**
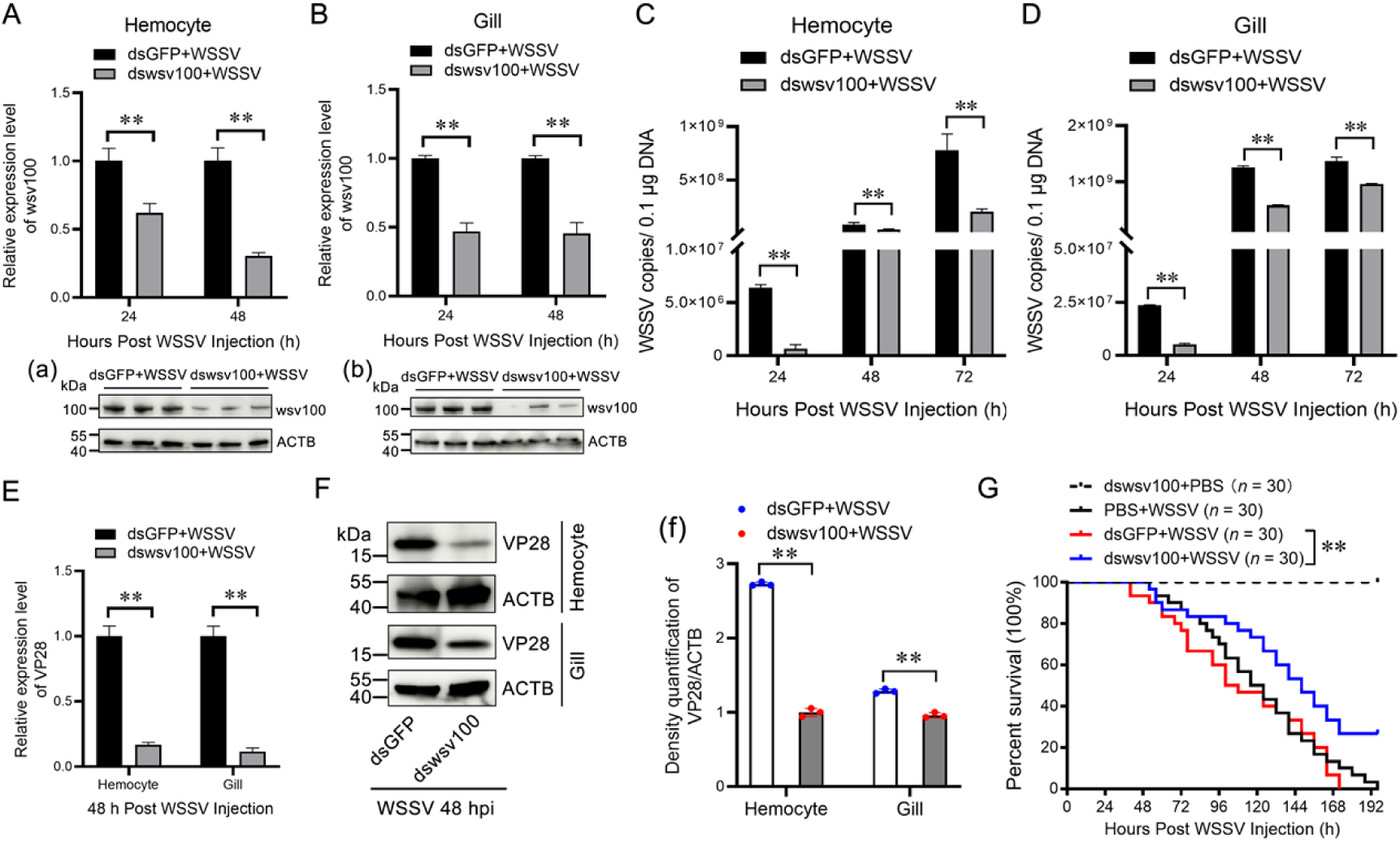
*wsv100* knockdown suppresses WSSV replication. (A-B) Silencing efficiency of *wsv100* in hemocytes (A) and gills (B) of WSSV-infected shrimp, assessed by qPCR. Protein levels of *wsv100* were analyzed by western blotting in hemocytes (panel a) and gills (panel b). Expression values were normalized against *EF-1α* and *β-Actin* using the Livak (2⁻^ΔΔCT^) method. Data are presented as mean ± SD from triplicate assays. (C-D) WSSV copy numbers in hemocytes (C) and gills (D) post-WSSV infection were quantified using absolute qPCR. (E) Transcriptional expression of the *VP28* gene was measured by qPCR in hemocytes and gills following WSSV infection. (F) VP28 protein levels were evaluated by western blotting in hemocytes and gills. (f) Quantification of VP28 protein expression from three independent repeats was performed using ImageJ. (G) Survival rates of WSSV-infected shrimp post-*wsv100* knockdown were recorded at 4-hour intervals and analyzed using Kaplan-Meier plots (log-rank χ² test). All experiments were conducted in triplicate and consistently yielded similar results. Statistical analyses were performed using Student’s *t*-test for panels (A-F) and Kaplan-Meier analysis for panel (G).

### 2.3 Overexpression of wsv100 Promotes WSSV Replication

To further investigate wsv100’s role, recombinant wsv100 (rwsv100) was co-injected with WSSV *in vivo*. Coomassie staining and western blotting confirmed successful expression and purification of rwsv100 (Figs. 3A, 3B). Shrimp co-injected with rwsv100 exhibited significantly reduced survival rates compared to controls injected with recombinant TRX protein (*p* < 0.05; Fig. 3C). Absolute quantitative PCR revealed higher viral titers in rwsv100-treated shrimp at 12, 24, 48, and 72 hpi (Fig. 3D). Additionally, rwsv100-treated shrimp showed increased levels of wsv100 and VP28 proteins at 24 and 48 hpi, respectively (Figs. 3E, 3e, 3F, 3f). These results indicate that rwsv100 promotes WSSV replication, exacerbating the infection.

**Figure 3.**
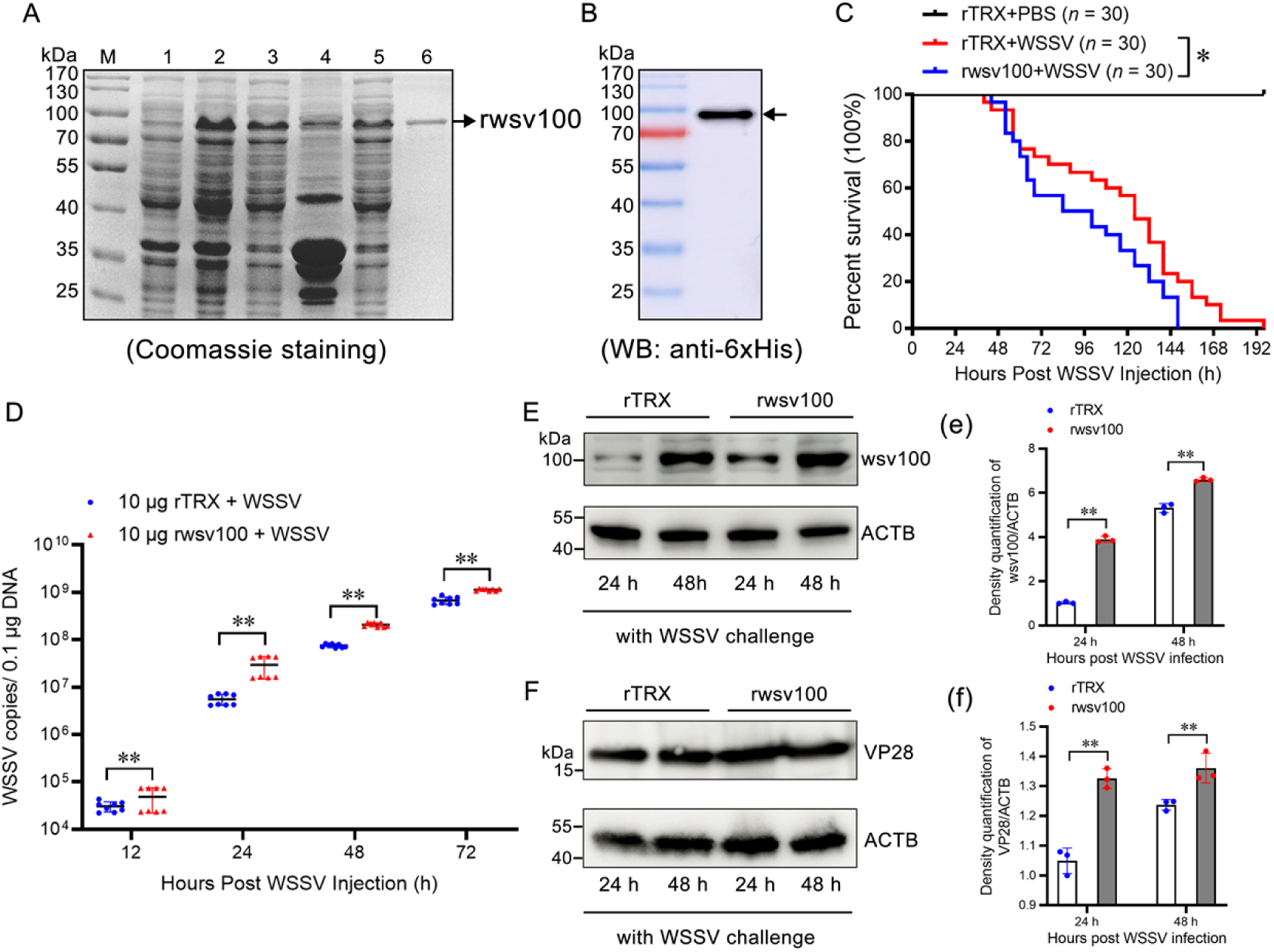
Overexpression of wsv100 promotes WSSV replication. (A) Recombinant expression and purification of rwsv100 in *E. coli*. Lane 1: uninduced *E. coli* transformed with rwsv100; Lane 2: induced *E. coli* expressing rwsv100; Lane 3: supernatant of ultrasonic lysate from *E. coli* expressing rwsv100; Lane 4: precipitate of lysed *E. coli*; Lane 5: supernatant after binding to His-tagged resin; Lane 6: purified rwsv100 protein (indicated by a black arrow). (B) Western blot analysis of purified rwsv100 protein using an anti-6 × His antibody (indicated by a black arrow). (C) Survival rates of shrimp injected with 10 μg rwsv100or control TRX, combined with WSSV inoculum. Survival rates were recorded every 4 hours and analyzed using a Kaplan-Meier plot (log-rank χ² test). (D) Quantification of WSSV copy numbers in shrimp gills following 10 μg rwsv100 application, measured *in vivo* using absolute qPCR. (E-F) Protein expression levels of rwsv100 (E) and VP28 (F) in shrimp gills post-*rwsv100* application at 24 and 48 hours post-infection (hpi), assessed by western blotting. (e-f) Quantification of rwsv100 (e) and VP28 (f) protein expression levels were performed using ImageJ. Statistical significance was determined using Student’s *t*-test (**p < 0.01, \**p* < 0.05). All experiments were conducted in triplicate and yielded consistent results.

### 2.4 Interaction Between wsv100 and Dorsal/IMD

To elucidate the mechanism by which WSSV utilizes wsv100 to evade the shrimp innate immune response, we performed His-tag pulldown assays using recombinant wsv100 (rwsv100) to isolate potential interacting proteins from hemocytes of WSSV-infected shrimp. Protein bands that were distinct from control TRX pull-downs were excised and analyzed using LC-MS/MS (Fig. 4A). The analysis identified several candidate proteins, including DnaJ-like protein, hemocyanin, C-type lectin, MAPK, tubulin, Pellino, actin, Dorsal, death domain-containing protein, and 14-3-3 protein (Fig. 4B). Proteins matching two or more unique peptides were classified as high-confidence interactors. Among these candidates, Dorsal and the death domain-containing protein stood out as high-confidence interactors with multiple peptide matches (Supplementary Figs. 2A, 2B). Notably, Dorsal is a crucial transcription factor in the canonical Toll pathway, responsible for inducing antimicrobial peptides to combat viral invasion [18]. Additionally, shrimp IMD, a death domain-containing protein, plays a key role as an adaptor protein in the IMD pathway, contributing to antiviral immune responses [12].

**Figure 4.**
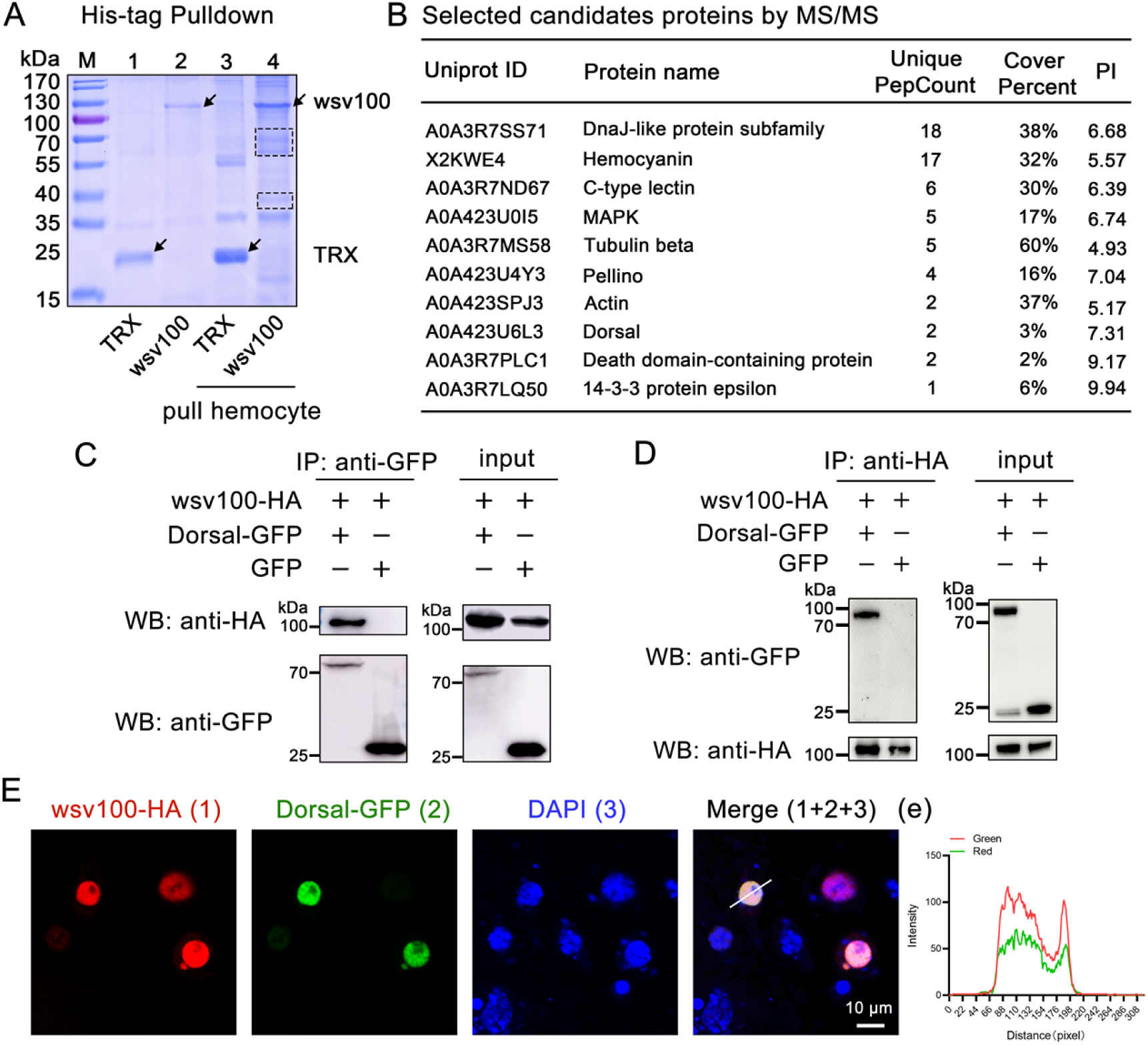
wsv100 interacts with Dorsal and IMD. (A) Pull-down assay of wsv100 in hemocyte lysates. Recombinant wsv100 protein was used to capture interacting proteins from hemocyte lysates of shrimp via His-tag pull-down assays. Protein bands corresponding to the wsv100 pull-down were excised and analyzed by liquid chromatography-tandem mass spectrometry (LC-MS/MS). (B) Proteomic identification of potential wsv100-interacting proteins based on LC-MS/MS analysis. (C-D) Co-immunoprecipitation (CoIP) assay to confirm interactions between wsv100 and Dorsal or IMD. Proteins were expressed in S2 cells, and immunoprecipitation was performed using anti-GFP antibodies (C) or anti-HA antibodies (D). Immunoprecipitates were detected with corresponding secondary antibodies. (E) Colocalization of wsv100 with Dorsal in *Drosophila* S2 cells. wsv100 was detected using mouse anti-HA antibodies followed by Alexa Fluor 594-conjugated secondary antibodies. Nuclei were stained with DAPI. Scale bar = 10 μm. (e) Quantitative analysis of fluorescence colocalization. The colocalization intensity of wsv100 with Dorsal was analyzed, with complete colocalization indicated by overlapping fluorescence peaks and maxima shifts of less than 20 nm. All experiments were performed in triplicate and consistently yielded similar results.

To confirm these interactions, we conducted co-immunoprecipitation (Co-IP) assays to examine the interaction between wsv100 and Dorsal or IMD. HA-tagged wsv100 and GFP-tagged Dorsal were co-expressed in S2 cells for this purpose. Using anti-GFP magnetic beads to immunoprecipitate Dorsal, wsv100 was detected as a co-precipitant (Fig. 4C). Conversely, Dorsal were identified in the immunoprecipitates of wsv100 using anti-HA magnetic beads (Fig. 4D). Confocal microscopy further validated the colocalization of wsv100 with Dorsal in *Drosophila* S2 cells (Figs. 4E, 4e). Similarly, wsv100 was shown to bind IMD through Co-IP assays with both anti-GFP and anti-HA magnetic beads (Supplementary Figs. 3A, 3B). Confocal microscopy also confirmed the colocalization of wsv100 with IMD in S2 cells (Supplementary Figs. 3C, 3c).

To further delineate the interaction domains of wsv100, Dorsal, and IMD, we segmented wsv100 into five distinct regions (1–124 aa, 125–248 aa, 249–372 aa, 373– 496 aa, and 497–624 aa; Fig. 5A). Dorsal was segmented into three regions based on its RHD and IPT domains (S1: 1–78 aa, S2: 79–249 aa, S3: 250–400 aa; Fig. 5B), while IMD was divided into two regions based on its death domain (IMD-N: 1–81 aa, IMD-C: 82–180 aa; Supplementary Fig. 4A). Co-IP assays indicated that residues 125–248 of wsv100 interact with Dorsal (Figs. 5C, 5D), while residues 1–124, 249–372, 373– 496, and 497–624 of wsv100 interact with IMD (Supplementary Figs. 4B, 4C). Additionally, the RHD domain (79–249 aa) of Dorsal was identified as responsible for binding wsv100 (Figs. 5E, 5F), whereas the N-terminal region (1–81 aa) of IMD mediated its interaction with wsv100 (Supplementary Figs. 4D, 4E).

**Figure 5.**
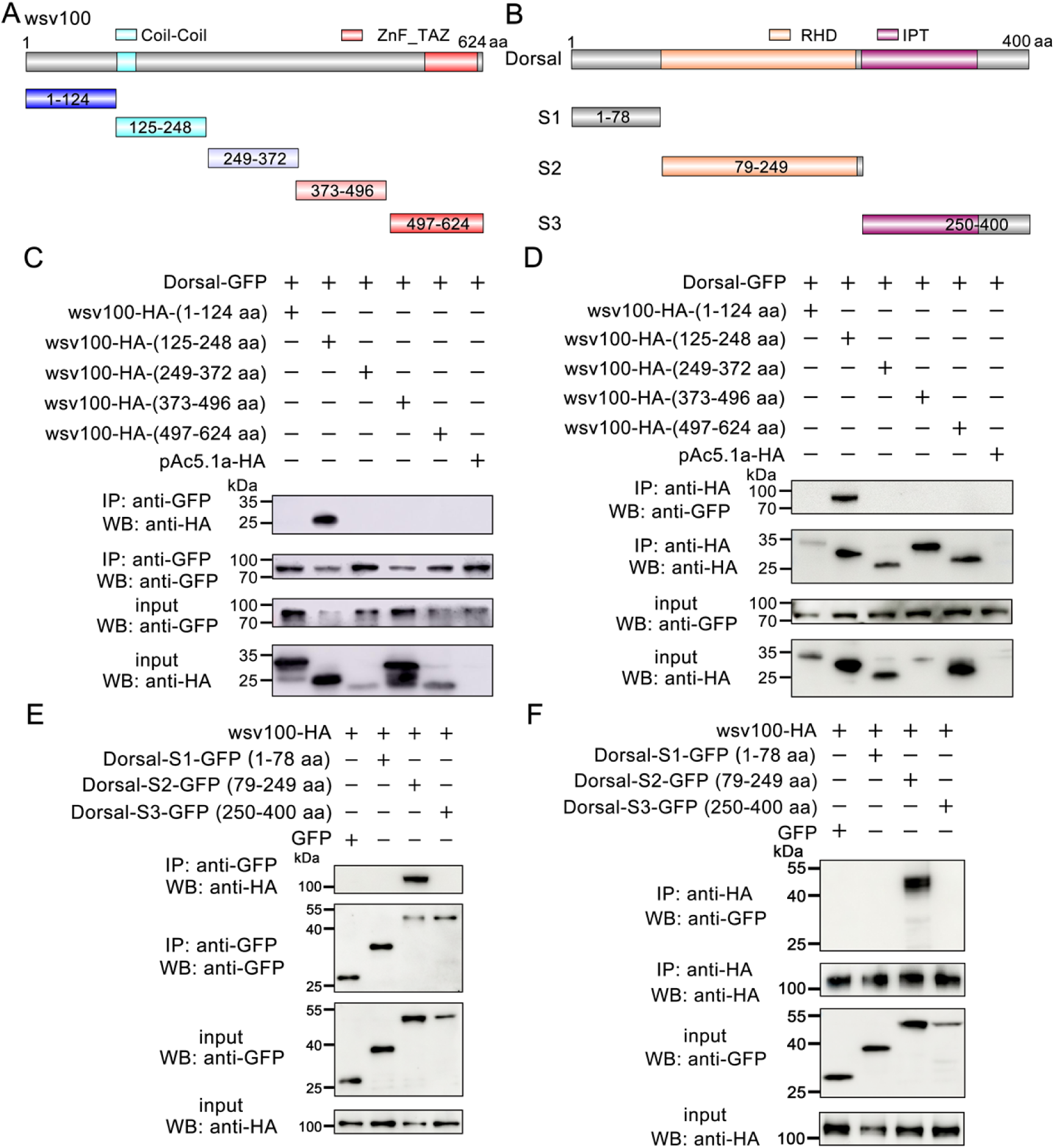
Domain mapping of wsv100-Dorsal associations. (A) Schematic representation of wsv100 and its truncation mutants used for domain mapping. (B) Schematic representation of Dorsal and its truncation mutants used for domain mapping. (C-D) Identification of the wsv100 domain interacting with Dorsal. GFP-tagged full-length Dorsal expressed in *Drosophila* S2 cells was immunoprecipitated, and HA-tagged wsv100 truncation mutants were detected in GFP-immunoprecipitates (C) or HA-immunoprecipitates (D). Input samples were analyzed using anti-GFP and anti-HA antibodies. (E-F) Identification of the Dorsal domain interacting with wsv100. HA-tagged full-length wsv100 expressed in *Drosophila* S2 cells was immunoprecipitated, and GFP-tagged Dorsal truncation mutants were detected in GFP-immunoprecipitates (E) or HA-immunoprecipitates (F). Input samples were analyzed using anti-GFP and anti-HA antibodies. All experiments were conducted in triplicate, yielding consistent and reproducible results.

Together, these results conclusively demonstrate that wsv100 interacts with Dorsal and IMD, targeting multiple components of the shrimp NF-κB signaling pathways.

### 2.5 wsv100 Inhibits Dorsal Phosphorylation and Nuclear Translocation

Phosphorylation and nuclear translocation of Dorsal is critical for its activation in the NF-κB pathway [18, 31]. Immunofluorescence experiments showed enhanced nuclear translocation of Dorsal in wsv100-knockdown shrimp (Figs. 6A, 6a). Western blotting revealed increased Dorsal phosphorylation in wsv100-silenced shrimp compared to controls (Figs. 6B, 6b). This was accompanied by increased nuclear localization of Dorsal and elevated expression of eight Dorsal-regulated AMPs (*anti-lipopolysaccharide factors* ALF1–4 and *lysozymes* LYZ1–4) in gill tissues (Figs. 6C–E). These results suggest that wsv100 inhibits Dorsal phosphorylation, thereby suppressing AMP production and enabling immune evasion.

**Figure 6.**
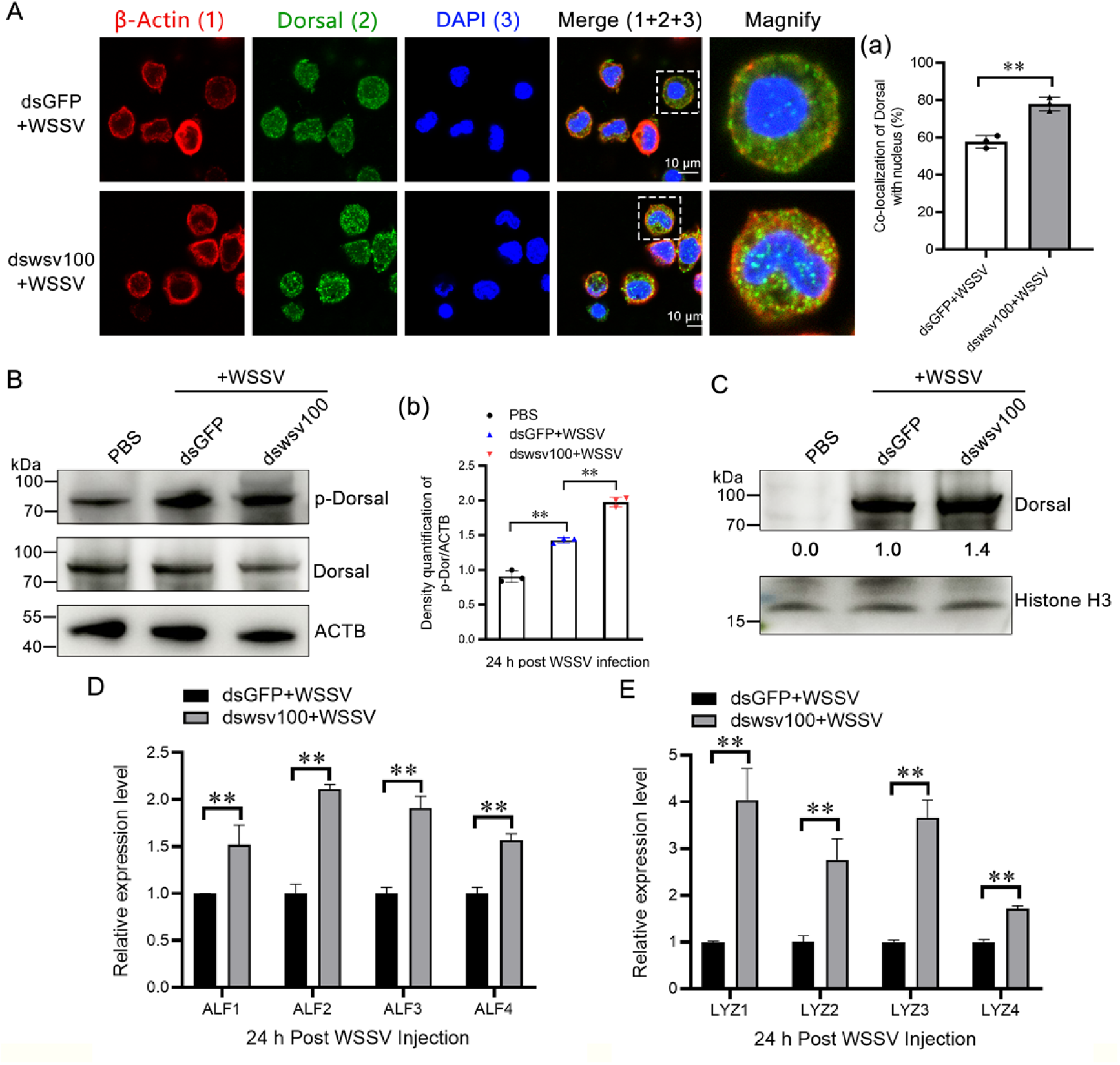
wsv100 inhibits the phosphorylation of Dorsal. (A) Effect of wsv100 knockdown on Dorsal nuclear translocation following WSSV infection. Hemocytes were collected 24 hours post-infection and analyzed via immunocytochemistry using an anti-Dorsal antibody. Scale bar = 10 μm. (a) Quantitative analysis of Dorsal co-localization with Hoechst-stained nuclei in hemocytes was performed using ImageJ software. (B) Phosphorylation of Dorsal in WSSV-infected shrimp after *wsv100* knockdown. Western blot analysis was conducted using an anti-Dorsal (phospho S276) antibody. (b) Quantitative analysis of Dorsal phosphorylation was performed using ImageJ software based on three independent experiments. (C) Nuclear distribution of Dorsal in WSSV-infected shrimp after *wsv100* knockdown. Western blotting was performed using an anti-Dorsal antibody to evaluate nuclear localization. (D-E) Expression levels of AMPs, including ALF1-4 (D) and LYZ1-4 (E), post-WSSV infection were analyzed in *wsv100* knockdown shrimp. Statistical significance of differences in expression levels was determined using Student’s *t*-test (\*\**p* < 0.01; \**p* < 0.05). All experiments were performed in triplicate, yielding consistent and reproducible results.

### 2.6 wsv100 Competes with Pelle for Dorsal Binding, Inhibiting Dorsal Phosphorylation and Antiviral Activity

In *Drosophila*, Dorsal phosphorylation is mediated by the kinase Pelle, which enhances Dorsal’s gene regulatory activity after its release from Cactus [10, 32]. In this study, the interaction between LvPelle and the RHD domain of LvDorsal was confirmed through GST pull-down assays performed *in vitro*. Recombinant GST-Pelle-N, GST-Pelle-C, and GST beads were co-incubated overnight at 4 °C with His-tagged Dorsal-RHD. GST alone was used as a negative control. The results demonstrated that both GST-Pelle-N and GST-Pelle-C interact with Dorsal-RHD (Fig. 7A; Supplementary Figs. 5A, 5B, 5C). This interaction was further validated using His-tagged pull-down assays (Fig. 7B).

**Figure 7.**
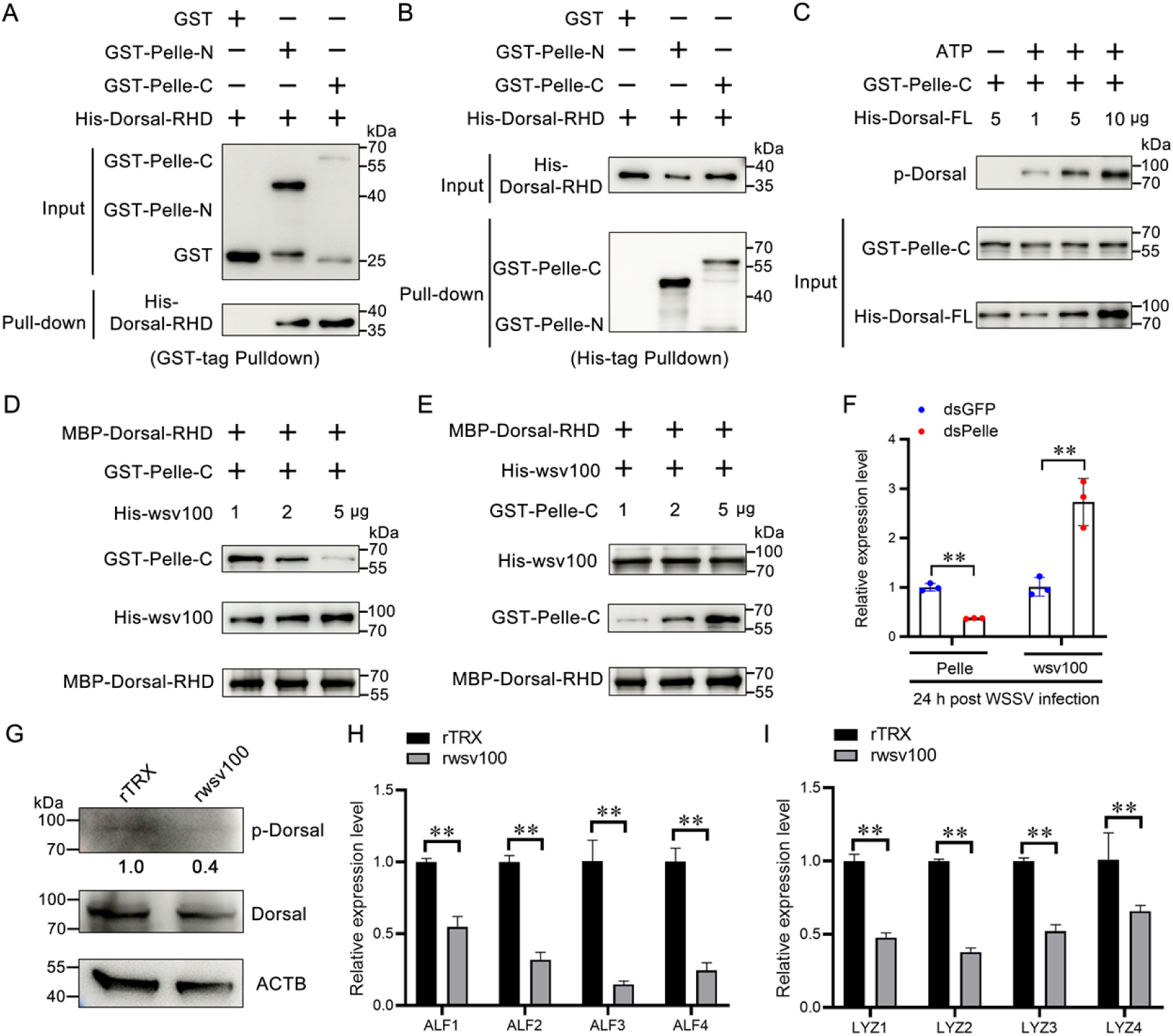
wsv100 competes with Pelle for Dorsal binding. (A-B) Interaction between Dorsal-RHD and Pelle domains was verified by in vitro pull-down assays. (A) GST-tagged pull-down assay using GST or recombinant TRX (rTRX) as controls. (B) His-tagged pull-down assay confirming the interaction. (C) Phosphorylation of Dorsal-FL protein by Pelle kinase was analyzed in vitro using ATP and kinase buffer. Identical protein inputs were confirmed by western blotting. (D) Binding ability of Pelle-C to Dorsal-RHD in the presence of increasing concentrations of wsv100 protein. (E) Binding ability of wsv100 protein to Dorsal-RHD in the presence of increasing concentrations of Pelle-C. (F) Effect of Pelle knockdown on *wsv100* expression in WSSV-infected shrimp, analyzed by qPCR. (G) Dorsal phosphorylation levels in rTRX- and rwsv100-injected shrimp were evaluated using western blotting with an anti-Dorsal (phospho S276) antibody. (H-I) Expression of AMPs, including ALF1-4 (H) and LYZ1-4 (I), in shrimp injected with rwsv100 protein, analyzed by qPCR. Statistical significance was determined using Student’s *t*-test (\*\**p* < 0.01; \**p* < 0.05). All experiments were conducted in triplicate and yielded consistent results.

The C-terminal kinase domain of Pelle (Pelle-C) is responsible for phosphorylating substrates. *In vitro* phosphorylation assays revealed that increasing concentrations of LvDorsal-FL protein led to a steady increase in its phosphorylation by LvPelle-C (Fig. 7C; Supplementary Fig. 5D). To investigate the competitive binding between wsv100 and Pelle to LvDorsal, a competitive assay was conducted. The results showed that increasing concentrations of His-tagged wsv100 protein reduced the binding of MBP-Dorsal-RHD to GST-Pelle-C (Fig. 7D; Supplementary Fig. 5E). However, the binding of His-wsv100 to MBP-Dorsal-RHD remained unaffected by increasing concentrations of GST-Pelle-C (Fig. 7E). Additionally, knockdown of LvPelle in shrimp significantly increased the transcription of wsv100 (Fig. 7F). Notably, injection of purified wsv100 protein into shrimp led to a marked reduction in LvDorsal phosphorylation in hemocytes compared to the TRX control group (Fig. 7G). Similarly, injection of wsv100 protein resulted in a significant decrease in the expression levels of AMPs, including ALF1-4 and LYZ1-4 (Figs. 7H, 7I), mirroring the effects observed with reduced LvDorsal activity. These findings collectively suggest that the WSSV wsv100 protein competes with LvPelle for binding to LvDorsal, thereby inhibiting Dorsal phosphorylation.

## 3. Discussion

The evolutionary arms race between viruses and their hosts drives the refinement of host immune defenses and the development of sophisticated viral counterstrategies [2]. The NF-κB pathway, traditionally recognized for its antibacterial role in invertebrates, is increasingly understood as a critical component of antiviral immunity [33]. This study provides novel insights into how WSSV subverts the shrimp NF-κB signaling pathway. By directly targeting the core components Dorsal and IMD, the viral protein wsv100 inhibits Dorsal phosphorylation and nuclear translocation, suppressing the expression of antimicrobial peptides (AMPs) and enabling immune evasion. These findings underscore the central role of NF-κB signaling in antiviral defense and highlight its vulnerability to viral exploitation.

Dorsal, a key transcription factor in the Toll pathway, is essential for immunity against fungi and Gram-positive bacteria [31]. However, accumulating evidence reveals its antiviral role across species. In *Drosophila*, Dorsal-deficient mutants show increased susceptibility to *Drosophila C virus* (DCV), while in shrimp, Dorsal knockdown enhances WSSV replication [18, 20, 34]. Viruses, in turn, have evolved diverse strategies to target NF-κB pathways: for instance, Rice stripe virus (RSV) NS4 protein inhibits NF-κB by competing with MSK2 kinase for Dorsal binding, and HSV-1 UL24 can suppress the transcription of NF-κB via direct binding to p65 [35, 36]. Additionally, In *Drosophila*, mutations of several IMD pathway genes displayed increased sensitivity to Cricket Paralysis virus (CrPV) infection and high virus loads, and *Drosophila* IMD pathway mediates the antiviral response to alphaviruses [13, 37]. Our study reveals a distinct mechanism: wsv100 directly interacts with Dorsal and IMD, preventing Dorsal phosphorylation by Pelle kinase and nuclear translocation. This direct interaction highlights a novel viral strategy to hijack a critical immune pathway.

Phosphorylation of Dorsal is a pivotal step in NF-κB activation, enabling its nuclear translocation and binding to target gene promoters [18]. In shrimp, the conserved serine 342 of LvDorsal, analogous to RPSD motifs in vertebrate p65, is critical for its antiviral activity [20]. This study demonstrates that *wsv100* competes with Pelle, a Toll pathway kinase, for binding to Dorsal, thereby blocking its phosphorylation. Notably, this mirrors the mechanism employed by RSV NS4 protein in plant arbovirus, where competitive inhibition of Dorsal phosphorylation suppresses host defenses [35]. Such parallels between invertebrate and plant systems underscore the evolutionary conservation of NF-κB as a target of viral immune evasion. Our findings also suggest that the NF-κB pathway, traditionally viewed as antibacterial in invertebrates, plays a broader antiviral role. By encoding wsv100 to disrupt this pathway, WSSV emphasizes the pathway’s importance in antiviral defense and its susceptibility to exploitation. The dual role of NF-κB in antibacterial and antiviral immunity likely represents a key evolutionary battleground between hosts and pathogens.

Both the Toll and IMD pathway activate NF-κB transcription factors, Dorsal and Relish, to regulate antimicrobial peptides (AMPs) production [38]. In shrimp, Dorsal binds to AMP gene promoters such as those of ALF1 and LYZ1 [20], and our study shows that wsv100 inhibits this regulation, suppressing AMPs expression. AMPs like ALFPm3, LvBigPEN and defensin play a direct role in WSSV defense by destabilizing viral particles, highlighting their importance in shrimp immunity [39–41]. Notably, the NF-kB transcription factors (Dorsal and Relish) participated in the regulation of BigPEN and LvDBD expression [40, 41]. Beyond AMPs, Dorsal likely regulates a broader network of immune effectors. For example, in *Drosophila*, DmDorsal modulates over 40 target genes, including autophagy-related genes [42]. Similarly, RSV NS4 protein inhibits LsDorsal phosphorylation, reducing immune effector expression such as LsZN708 [35]. Future studies should investigate whether *wsv100* also affects other immune-related genes downstream of LvDorsal, including those involved in autophagy or stress responses.

From an applied perspective, these findings offer potential avenues for developing antiviral strategies. Identifying small molecules or genetic tools to disrupt the wsv100-Dorsal interaction could help restore NF-κB pathway activity and enhance shrimp immunity. Additionally, understanding whether other WSSV proteins cooperate with wsv100 to modulate NF-κB could provide further targets for intervention. Beyond shrimp aquaculture, the insights gained here may extend to other aquatic pathogens, offering a framework for investigating viral immune evasion strategies in diverse invertebrate systems. While this finding establishes the critical role of wsv100 in inhibiting NF-κB signaling, some questions remain. For instance, how does the interaction between wsv100 and IMD affect the IMD pathway’s broader role in antiviral defense, which will be essential for a comprehensive understanding of WSSV immune evasion strategies.

In conclusion, herein, we uncover a novel mechanism by which WSSV antagonizes the shrimp NF-κB signaling pathway. Specifically, the IE gene wsv100 competitively binds to Dorsal, preventing its phosphorylation by Pelle kinase and nuclear translocation (Fig. 8). This immune evasion strategy suppresses AMP production, facilitating viral infection (Fig. 8). These findings advance our understanding of host-virus interactions, highlighting the NF-κB pathway as a critical target of viral exploitation and a potential focus for antiviral interventions.

**Figure 8.**
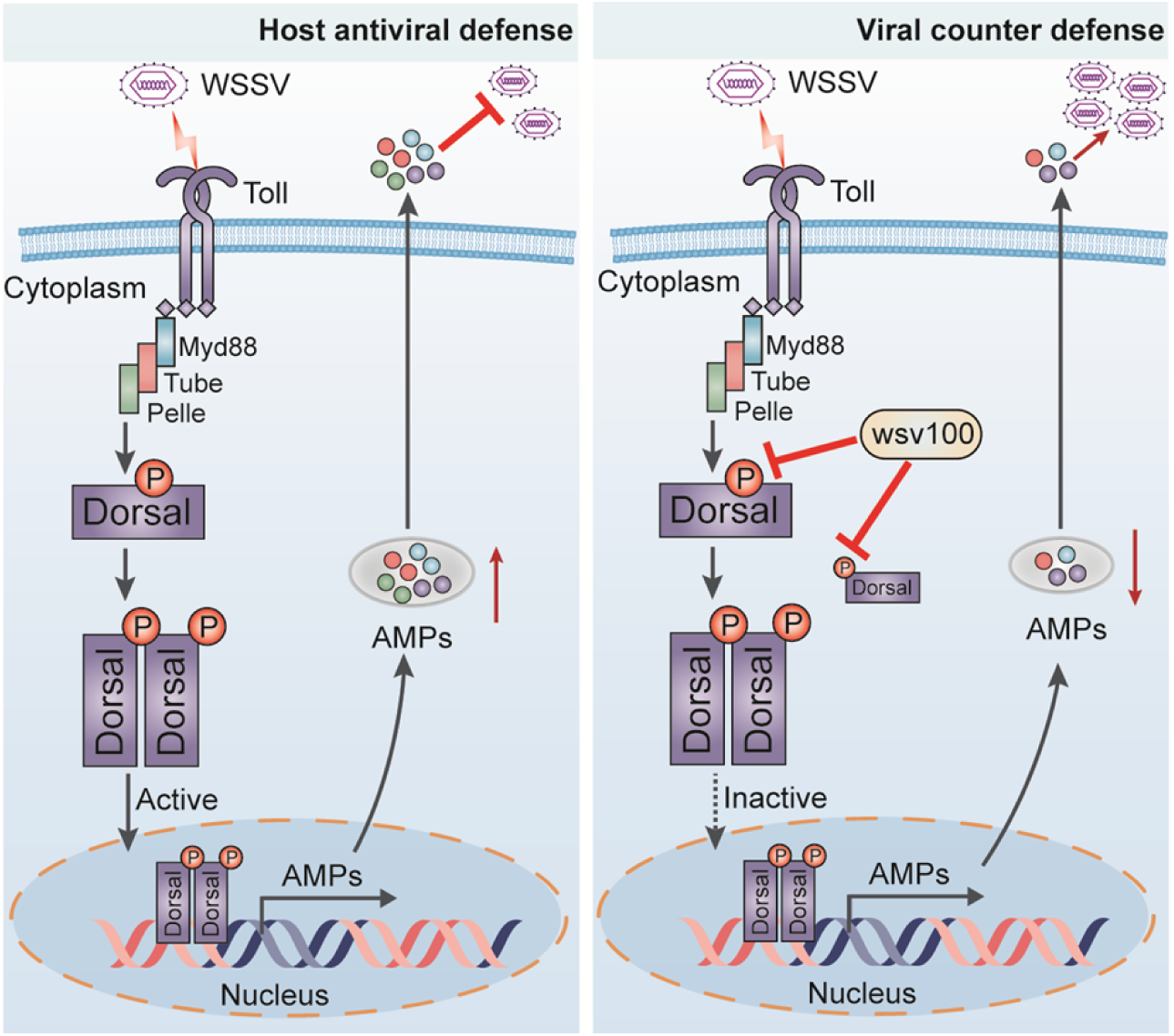
Proposed model for the mechanism by which *wsv100* evades NF-κB pathway-mediated immune responses in shrimp. The Toll receptor in shrimp recognizes WSSV infection, leading to the activation of the NF-κB pathway. In this pathway, the transcription factor Dorsal is phosphorylated by the kinase Pelle and translocates to the nucleus, where it induces the expression of downstream immune effectors, such as antimicrobial peptides (AMPs), to defend against WSSV infection (left). However, WSSV has evolved an antidefense strategy through its immediate-early (IE) protein *wsv100*, which antagonizes NF-κB activity by competitively binding to Dorsal and preventing its interaction with Pelle. This interaction suppresses Dorsal phosphorylation and nuclear translocation, thereby inhibiting the immune response and facilitating viral infection (right).

## 4. Materials and methods Animal and pathogens

Healthy specimens of *Litopenaeus vannamei* (approximately 5 g each), were used for our experiments. These specific pathogen-free (SPF) shrimp were sourced from Hisenor Company’s shrimp farm in Maoming, Guangdong Province, China. The shrimp were maintained in a recirculating water tank system filled with air-pumped seawater at a salinity of 25‰ and a constant temperature of 25 ℃. They were fed thrice daily to satiation with a commercial shrimp diet provided by HAID Group, Guangzhou, China. The inoculum for the WSSV, Chinese strain (AF332093), was prepared following previously described methodologies [43].

WSSV was extracted from the muscle tissue of WSSV-infected shrimp and subsequently stored at –80 ℃. Prior to injection, the muscle tissue was homogenized to prepare the WSSV inoculum, containing approximately 1 × 10^5 copies in 50 μl of Phosphate-Buffered Saline (PBS; 140 mM NaCl, 2.7 mM KCl, 10 mM Na_2_HPO_4_, 1.8 mM KH_2_PO_4_, pH 7.4). For the pathogenic challenge experiments, each shrimp was administered an intraperitoneal injection of 50 μl of the WSSV solution into the second abdominal segment. Injections were performed using a 1-ml syringe from Becton Dickinson, Shanghai, China.

### RNA extraction, cDNA synthesis, and DNA and protein extraction

Total RNA was extracted from different tissues of shrimp using the Eastep Super Total RNA Extraction Kit (Promega, Shanghai, China). The genomic DNA of shrimp tissues was extracted using a genomic DNA extraction kit (Vazyme, Nanjing, China). First-strand cDNA synthesis was performed using a cDNA synthesis kit (Accurate Biology, Hunan, China), according to the manufacturer’s instructions. Protein samples from gills and hemocytes were homogenized separately in IP lysis buffer (25 mM Tris-HCl pH 7.4, 150 mM NaCl, 1 mM EDTA, 1% NP-40, 5% glycerin; Thermo Scientific) with a protease and phosphatase inhibitor cocktail (Merck, Kenilworth, NJ, USA), and then centrifuged at 16,000 × *g* for 10 minutes at 4 ℃ to collect the supernatant for further test.

### Quantitative PCR (qPCR)

qPCR assays were performed to assess the transcription levels of wsv100 in WSSV infection or the *in vivo* RNAi experiments. The expression of wsv100 (GenBank accession No. AWQ63223) was detected using LightCycler480 System (Roche, Basel, Germany) in a final reaction volume of 10 μl, which was comprised of 1 μl of 1: 10 cDNA diluted with ddH_2_O, 5 μl of SYBR Green Pro Taq HS Mix (Accurate Biology, Hunan, China) and 250 nM of specific primers (Supplementary table 1).

For WSSV stimulation, the treated group were injected with 50 μl WSSV (∼1 × 10^5^ copies), and the control group was injected with PBS solution. Hemocyte and gill of challenged shrimps were collected at 0, 4, 8, 12, 24, 36, 48, and 72 h after injection. The expression of each gene was calculated using the Livak (2^-ΔΔCT^) method after normalization to *EF-1α* (GenBank accession No. GU136229) and *β-Actin* (GenBank accession No. AF186250) as described previously [44].

The transcription of *ALF1* (GenBank accession No. AVP74301), *ALF2* (GenBank accession No. AVP74302), *ALF3* (GenBank accession No. ABB22831), *ALF4* (GenBank accession No. AHG99284), *LYZ1* (GenBank accession No. ABD65298), *LYZ2* (GenBank accession No. AAL23948), *LYZ3* (GenBank accession No. MH357347), *LYZ4* (GenBank accession No. AVP74306), *VP28* (GenBank accession No. NP_477943.1) and *Pelle* (GenBank accession No. KC346864.1) were also detected by qPCR. Primer sequences are listed in Supplementary table 1.

### Detection of viral loads by absolute quantitative PCR (ab-qPCR)

To measure viral titers in shrimp, absolute quantitative PCR (ab-qPCR) was employed. This process utilized a set of primers, wsv069 (WSSV32678-F/WSSV32753-R), targeting a single-copy gene of the WSSV, along with a Taq-Man fluorogenic probe (WSSV32706), as detailed in previous studies [45]. Gill tissue samples were collected from WSSV-infected shrimp. DNA was extracted from these samples using the methods described above. The concentration of WSSV genome copies was quantified by ab-qPCR, using the specific primers WSSV32678-F/WSSV32753-R and the TaqMan fluorogenic probe, as listed in Supplementary table 1. To ensure accuracy, each shrimp sample underwent three replicates of ab-qPCR. The number of WSSV genome copies was calculated and normalized against 0.1 μg of shrimp tissue DNA.

### RNA interference (RNAi) for gene knockdown

The T7 RiboMAX Express RNAi System kit (Promega, Shanghai, China) was employed for the synthesis of double-stranded RNA (dsRNA) targeting GFP, wsv100 and Pelle. The specific primers used for dsRNA synthesis are listed in Supplementary table 1. The integrity and quality of the synthesized dsRNA were verified using 1.5% agarose gel electrophoresis and quantified with a NanoDrop 2000 spectrophotometer (Thermo Scientific, Shanghai, China). For the experimental treatments, shrimp were injected with 2 μg/g of body weight of the respective dsRNA, dissolved in 50 μl of PBS. Control groups received injections of GFP dsRNA and PBS. Shrimp tissues were collected from the shrimp 48 hours post-dsRNA injection. The efficiency of RNAi was evaluated by qPCR using the corresponding primers for each gene. This assessment aimed to confirm the knockdown of the targeted genes in the experimental groups compared to the controls as described previously [46].

### Survival assays

For wsv100 knockdown, healthy shrimp were divided into two groups (*n* = 30 each) and received an intramuscular injection of 10 μg dsRNA solution mixed with WSSV crude extract. The survival rates of each group were recorded at 4-hour intervals. The Mantel–Cox (log-rank χ^2^ test) method was subjected to analyze differences between groups using GraphPad Prism software (Graphpad, San Diego, CA, USA).

For rescue experiments, 10 μg rwsv100 was first incubated with WSSV crude extract, and then the mixture was inoculated into the experimental shrimp by injection. The TRX protein was used as a control. The survival rates of each group were recorded at 4-hour intervals. Differences in survival between groups were analyzed using the Mantel-Cox (log-rank χ^2^ test) method, employing GraphPad Prism software (GraphPad Software, La Jolla, CA, USA).

### Recombinant protein expression and purification

The coding sequences of wsv100 was amplified by PCR using corresponding primers (Supplementary table 1) and subcloned into pET-32a (+) plasmid (Merck Millipore, Darmstadt, Germany). After confirmed by sequencing, the recombinant plasmid was transferred into *E. coli* Rosetta (DE3) cells (TransGen Biotech, Beijing, China). Then, positive clones harboring the desired fragment were selected for inducing expression. After 12 hours of induction with 0.1 mM Isopropyl β-D-Thiogalactoside (IPTG) at 16 ℃, cells were pelleted by centrifugation and sonicated for 1 hour on ice water. The supernatant from the sonicated proteins was purified by using Ni-NTA agarose (Qiagen, Düsseldorf, Germany) according to the manufacturer’s instructions. The purity of the recombinant wsv100 (rwsv100) was assessed by Coomassie staining and Western blot analysis. To remove most of the endotoxin contamination, cold 0.1% Triton X-114 was used to fully wash the column before final elution of proteins with elution buffer [47]. The concentration of purified rwsv100 was determined using a bicinchoninic acid protein assay kit (Beyotime Biotechnology, Shanghai, China).

The recombinant protein of Dorsal-RHD-His, Dorsal-FL-His, Dorsal-RHD-MBP, Pelle-N-GST, and Pell-C-GST were purified in the same way with appropriate resin (GST resin or MBP resin).

### SDS-PAGE and western blotting

Western blotting was performed to evaluate the protein levels of wsv100 and VP28. And then the proteins were separated on 12.5% SDS-PAGE gels and then transferred to polyvinylidene difluoride (PVDF) membranes (Merck Millipore, Darmstadt, Germany). After blocking with 5% nonfat milk diluted in Tris Buffered Saline with Tween 20 (TBST) buffer (150 mM NaCl, 0.1% Tween-20, 50 mM Tris-HCl, pH 8.0) for 1 hour, and the membrane was incubated with appropriate Abs. The rabbit anti-wsv100 (customized from GL biochem), rabbit anti-VP28 (produced in Abmart), mouse anti-β-Actin (CST, catalog no. 3700S) were used. The PVDF membranes were washed for three times with TBST and then incubated with 1:5000 goat anti-rabbit IgG (H+ L) HRP or 1:5000 goat anti-mouse IgG (H+ L) HRP secondary antibody (Promega, Shanghai, China) for 1 hour. Membranes were developed using an enhanced chemiluminescent blotting substrate (Thermo Fisher Scientific, Waltham, MA, USA), and the chemiluminescent signal was detected using the 5200 Chemiluminescence Imaging System (Thermo Fisher Scientific, Waltham, MA, USA). For relative densitometry of VP28, Dorsal or p-Dorsal, the immunoblotted band volume was normalized to the corresponding internal protein volume in the lane, using the ImageJ software 1.6.0 (National Institutes of Health, Bethesda, MD). Statistical analysis of densitometry data from three independent experiments was performed by using the Student’s *t* test.

### Plasmid construction

The pAc5.1a-hemagglutinin (HA) and pAc5.1a-GFP plasmid was reconstructed by inserting a 3 × HA and GFP sequence at the N terminal of the multiple cloning site (MCS) of pAc5.1a-V5/His A (Thermo Scientific, Shanghai, China), respectively [48]. The wsv100, Dorsal, IMD, or their domain of fragment deletion mutants were cloned into the pAc5.1a-HA and pAc5.1a-GFP to generate wsv100-HA, wsv100-HA-(1-124 aa), wsv100-HA-(125-248 aa), wsv100-HA-(249-372 aa), wsv100-HA-(373-496 aa), wsv100-HA-(497-624 aa), Dorsal-GFP, Dorsal-GFP-S1, Dorsal-GFP-S2, Dorsal-GFP-S3, IMD-GFP, IMD-GFP-N, and IMD-GFP-C using corresponding primers (Supplementary table 1).

### Co-immunoprecipitation (Co-IP)

Co-immunoprecipitation (CoIP) assays *in vitro* were performed to confirm the interaction between proteins as described previously [49]. In brief, 48 h after transfection, *Drosophila* S2 cells were harvested and wash with ice-cold PBS three times and then lysed in IP lysis buffer (Thermo Scientific) with a protease and phosphatase inhibitor cocktail (Merck). The supernatants (400 μl) were incubated with 30 μl anti-HA magnetic beads (Thermo Scientifi) or anti-GFP magnetic beads (Smart-Lifesciences) at 4 ℃ for 4 h. Then the magnetic beads were washed with PBS for five time and subjected to SDS-PAGE assay. Five percent of each total cell lysate was also detected as the input control.

### GST pull-down assays and mass spectrometry (MS) analysis

To search for host proteins that interact with wsv100. The pull-down assay was conducted following established protocols with certain modifications [50]. We first incubated the purified His-tagged and wsv100-His tagged with Ni NTA-resin for 2 hours. Subsequently, this complex was further incubated with hemocyte lysate extracted from 20 shrimp for an additional 12 hours at 4 ℃. Post-incubation, the resin-bound proteins were washed six times using PBS containing 0.5 M NaCl and 1% Tween 20. The proteins were then eluted with 300 mM imidazole elution buffer (pH 8.0). The eluted proteins were separated by SDS-PAGE. Distinct bands on the gels were excised and submitted for liquid chromatography-tandem mass spectrometry analyses (LC-MS/MS) to Qinglianbio, Beijing, China.

To verify the interaction between Pelle and Dorsal, the pull-down assays were conducted. For GST pull-down assays, 5 μg rDorsal-RHD-His was incubated with 5 μg of GST-tagged Pelle-N or Pelle-C at 4 ℃ for 3 h by agitation. Subsequently, 20 μl of the GST resin were added and the agitation continued for another 2 h. The resin was washed four times with PBS. Finally, the resins were re-suspended in 50 μl of the SDS-PAGE sample buffer, boiled and analyzed by Western blot analysis using 6 × His antibody. The His pulldown assays were similar to the methods described above using Ni NTA resin.

### Nuclear protein extraction

Nuclear protein was extracted from dswsv100 application shrimp with WSSV infection using a NE-PER Nuclear and Cytoplasmic Extraction Reagents (Thermo Fisher Scientific). Specifically, hemocytes were collected into 1.5 ml tube. And the added CER buffer along with a protease inhibitor (Thermo Fisher Scientific). After incubation in an ice bath for 10 minutes, the supernatant was retained as the cytoplasmic protein. And then the remaining precipitate was treated with NER buffer, and the supernatant was retained as the nuclear protein following centrifugation at 16,000 × *g* for 10 minutes at 4 ℃. The extracted nuclear protein was then subject to Western blotting analysis using a rabbit anti-shrimp p-Dorsal antibody (Gene Create, China) reported in our previous study [20]. Anti-rabbit histone H3 antibody (CST, cat. no. 9715L) was used as nuclear reference antibody.

### Competitive binding assay

The competitive binding Assays were performed to investigate the competitive relationship between Pelle, wsv100, and Dorsal-RHD as described previously [35]. MBP-tagged Dorsal-RHD was incubated with MBP resin for 4 hours at 4 ℃. And the His-tagged wsv100 and GST-tagged Pelle-C protein with different amount were added with several different order for incubation at 4 ℃ for 12 hours. After centrifugation and wash four times with PBS. The protein bound to the beads were detected by Western blotting using antibodies specific to the His-tag, GST-tag and MBP-tag.

### Phosphorylation assay *in vitro*

To explore the relationship between Pelle kinase and LvDorsal, an phosphorylation assay was conducted *in vitro* as described previously [51]. In brief, the GST-tagged Pelle-C protein maintained at a constant amount and varying concentrations of His-Dorsal-FL protein with ATP (CST, cat. no. 9804S) were added into 10 × buffer (CST, cat. no. 9802S) to obtain a 1 × reaction system. In the control, the ATP was replaced by ddH_2_O. The mixtures were incubated for 1 hour at 30 ℃ on MIX Vertical and then boiled with 5 × loading buffer for Western blot analysis.

### Immunocytochemical staining

For analysis Dorsal or wsv100 translocation in hemocyte. Hemocyte from dsRNA application shrimp or WSSV infected, and the hemocyte were centrifuged at 500 × g for 10 min at 4 ℃. Then the cells were washed twice with PBS and spread onto a glass slide in 12-hole microtiter plates for 30 min. Subsequently, the glass slices in the wells were washed with PBS three times and fixed with 4% paraformaldehyde at 25 °C for 15 min. The hemocytes on the glass slides were washed with PBS three times and blocked with 3% bovine serum albumin (dissolved in PBS) for 60 min at 25 °C. The hemocyte was incubated with anti-Dorsal antibody (1: 300, diluted in blocking reagent) or anti-wsv100 (1: 300, diluted in blocking reagent) for 12 h at 4 °C. The hemocytes were then washed with PBS three times and then incubated with anti-rabbit IgG (H+L) Alexa Fluor 488 (CST, 1:1000 diluted in 2% BSA) for 60 min at 25 °C in the dark. The cell nuclei were stained with hoechst (Beyotime, Shanghai, China; cat. no. C1002) for 10 min. Finally, fluorescence was visualized on a confocal laser scanning microscope (Leica TCS-SP8, Wetzlar, Germany). We calculated the colocalization percentage of Dorsal with nucleus stained with hoechst, using the ImageJ according to the previously described methods [52].

To observe the co-localization of wsv100 with Dorsal and IMD. *Drosophila* S2 cells were seeded onto poly-L-lysine-treated glass cover slips in a 24-well plate with approximate 40% confluent. The S2 cells of each well were transfected with 0.5 μg pAc5.1-wsv100-HA, pAc5.1-Dorsal-GFP or pAc5.1-IMD-GFP using the MIK-3000 Transfection Reagent (MIKX, Shenzhen, China). Thirty-six hours post plasmid transfection, cell culture medium was removed, and cells washed with PBS three times. Cells were treated as described above with appropriate antibodies.

### Statistical analysis

All data were presented as means ± SD. Student’s *t* test was used to calculate the comparisons between groups of numerical data. For survival rates, data were subjected to statistical analysis using GraphPad Prism software (Graphpad, San Diego, CA, USA) to generate the Kaplan–Meier plot (log-rank χ^2^ test).

## Author contributions

Chaozheng Li, Jianguo He and Bang Xiao conceived and designed the experiments. Bang Xiao, Fang Kang, Qianqian Li, Junming Pan, Yue Wang and Chaozheng Li performed the experiments and analyzed data. Bang Xiao and Chaozheng Li wrote the draft manuscript. Bang Xiao, Chaozheng Li and Jianguo He acquired finding. Chaozheng Li was responsible for forming the hypothesis, project development, data coordination, and writing, finalizing, and submitting the manuscript. All authors discussed the results and approved the final version.

## Acknowledgement

This research was supported by National Natural Science Foundation of China (32373158/32441085/31930113), the open competition program of top ten critical priorities of Agricultural Science and Technology Innovation for the 14th Five-Year Plan of Guangdong Province (2022SDZG01), the Guangdong S&T programme (2024B1212060001), China Postdoctoral Science Foundation (2023M743987), and Guangdong Basic and Applied Basic Research Foundation (2023A1515110528). The funders had no role in study design, data collection and analysis, decision to publish, or preparation of the manuscript.

## Data availability

All reagents and experimental data are available within transparent methods or from corresponding author upon reasonable request.

## Ethics statement

All animal experiments were approved by Institutional Animal Care and Use Committee of Sun Yat-Sen University (Approval No. SYSU-IACUC-2024-B0640, 6 March 2024).

---

**Supplementary Figure 1.**
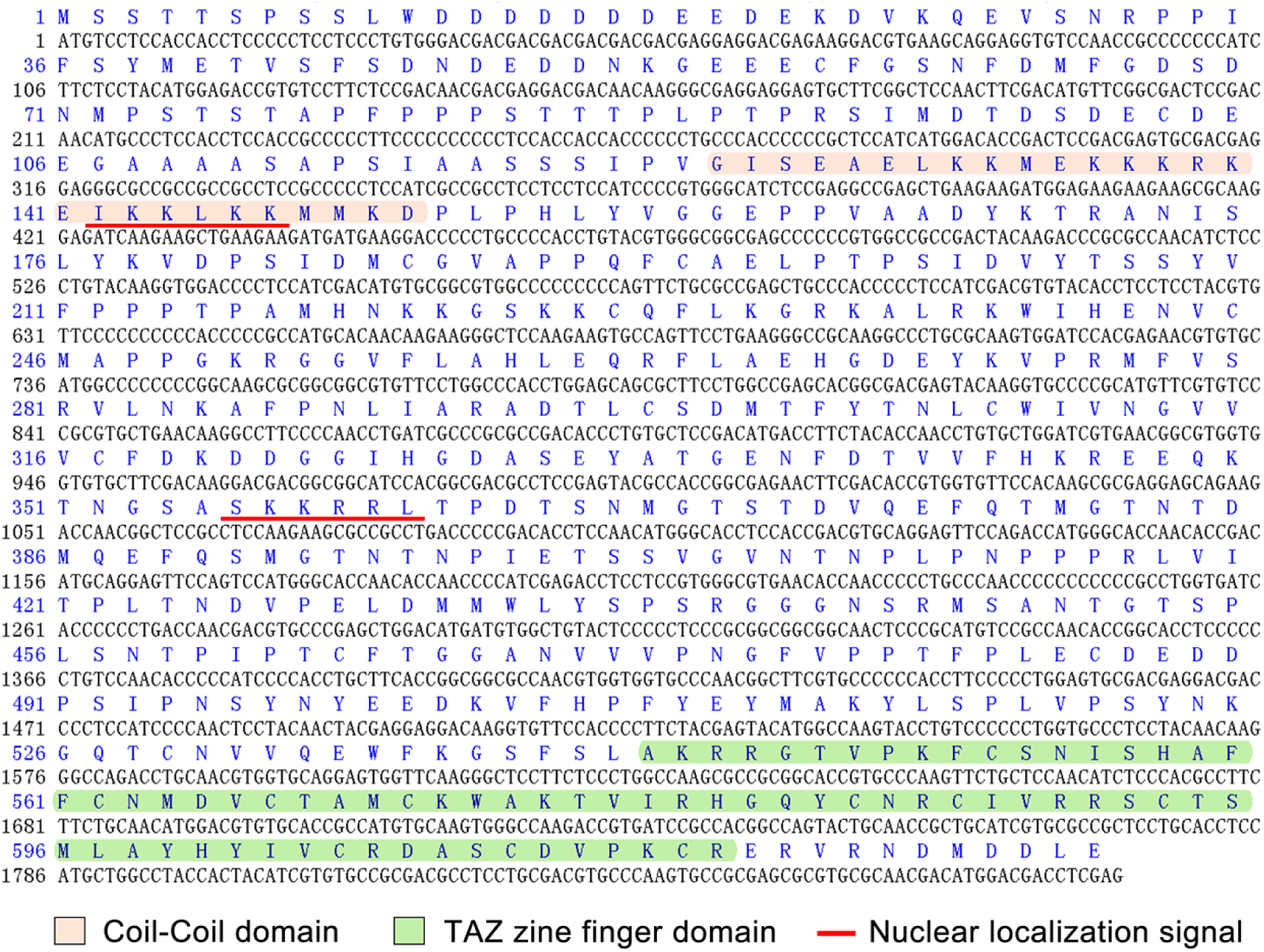
The amino acid sequence and conserved domain of wsv100. The orange represents coil-coil domain, the green represents TAZ zine finger domain, and the red underline represents nuclear localization signal.

**Supplementary Figure 2.**
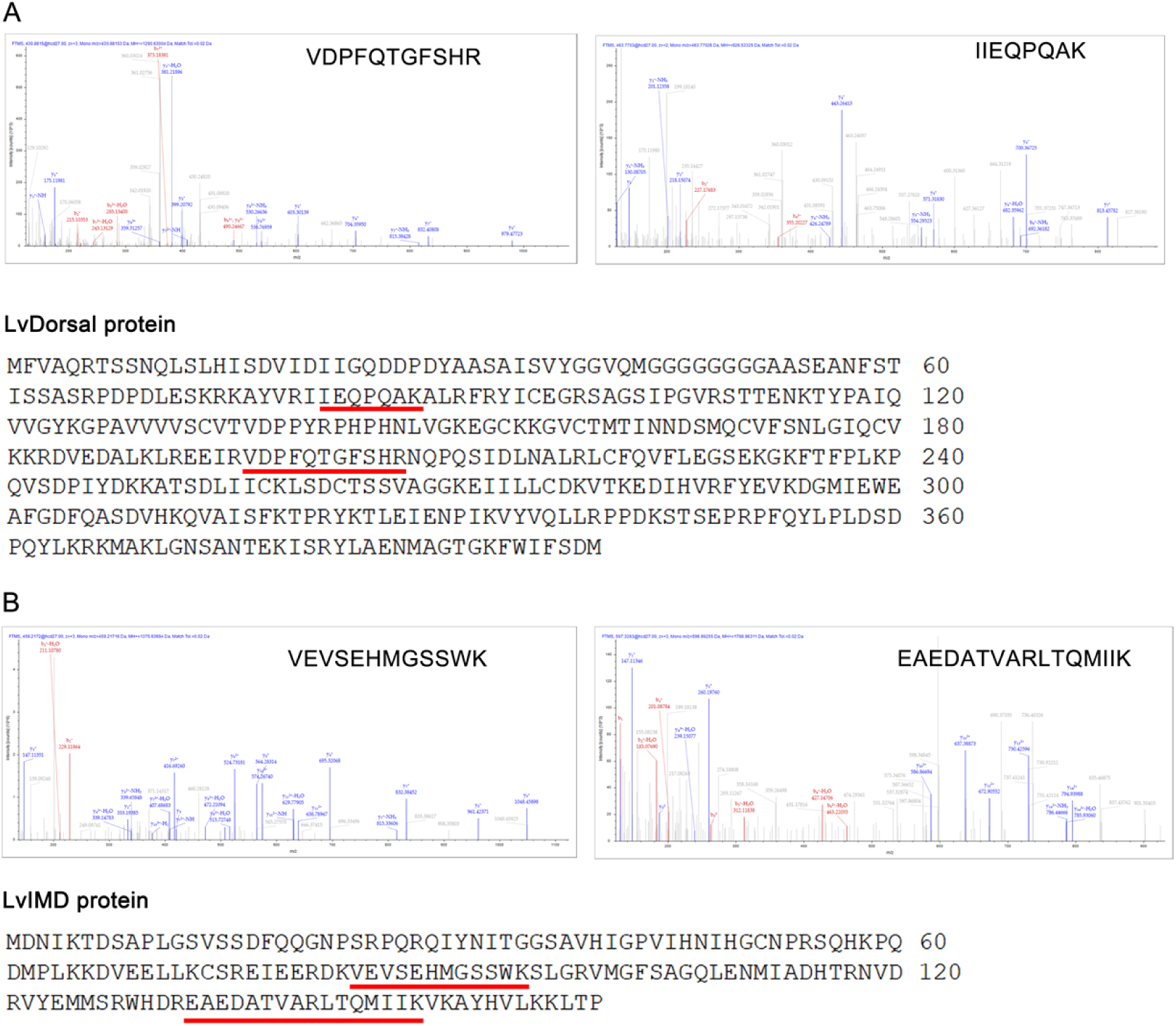
Identification of wsv100 interacting proteins via mass spectrometry. (A) MS/MS spectra was displayed for the identification of LvDorsal in the shrimp’s hemocyte lysates following His-tagged pulldown. Peptides identified through proteomic analysis are highlighted with underline, indicating their association with LvDorsal. (B) MS/MS spectra was displayed for the identification of LvIMD in the shrimp’s hemocyte lysates following His-tagged pulldown. Peptides identified through proteomic analysis are highlighted with underline, indicating their association with LvIMD.

**Supplementary Figure 3.**
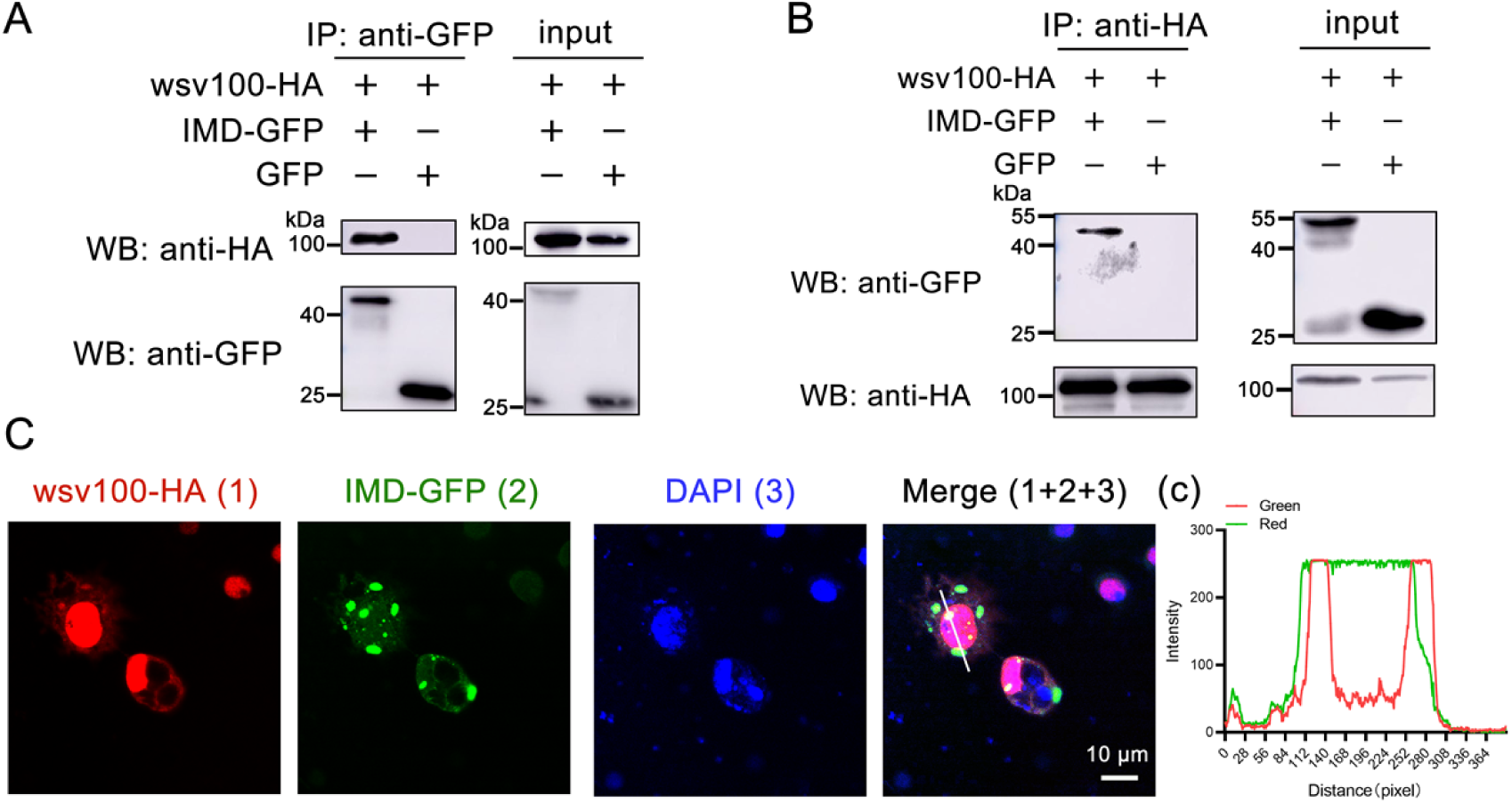
wsv100 interacts with IMD. (A, B) The interaction between wsv100 and IMD were performed using CoIP. Proteins were expressed in S2 cells, and IP was conducted using anti-HA (A) or anti-GFP antibodies (B). Immunoprecipitates were detected with corresponding secondary antibodies. (C) The colocalization of wsv100 with IMD in S2 cells. The wsv100 was detected with mouse anti-HA antibodies and anti-mouse Alexa Fluor 594. DAPI staining highlighted the nuclei. The scale bar represents 10 μm. (c) Quantitative analysis of fluorescence colocalization of wsv100 with IMD. Colocalization intensity was quantitatively analyzed, with complete colocalization indicated by overlapping peaks and maxima shifted by less than 20 nm. All experiments were representative of three biological replicates and yielded similar results.

**Supplementary Figure 4.**
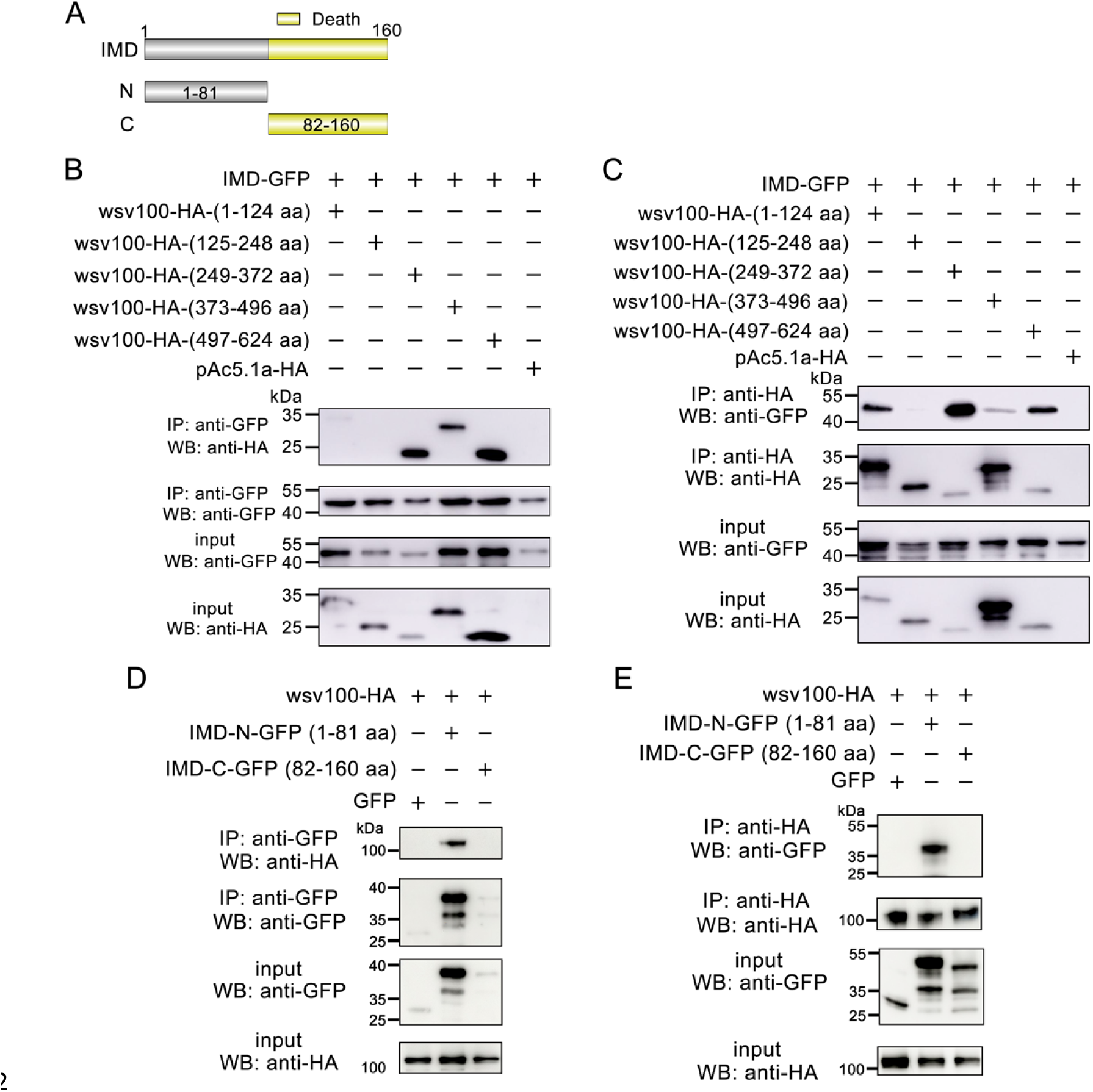
Domain mapping of wsv100-IMD associations. (A) Schematic diagram of IMD and its truncation mutants. (B-C) Identification of the wsv100 domain binding IMD. GFP-tagged IMD expressed in S2 cells were immunoprecipitated, and the presence of HA-tagged wsv100 truncation mutants were detected by GFP-immunoprecipitates (B) or HA-immunoprecipitates (C). Input samples were analyzed using anti-GFP and anti-HA antibodies. (D-E) Identification of the IMD domain binding to wsv100. HA-tagged wsv100 expressed in S2 cells was immunoprecipitated, and the presence of GFP-tagged IMD truncation mutants were detected by GFP-immunoprecipitates (D) or HA-immunoprecipitates (E). Input samples were analyzed using anti-GFP and anti-HA antibodies. Each experiment was conducted in triplicate, ensuring consistent and representative results.

**Supplementary Figure 5.**
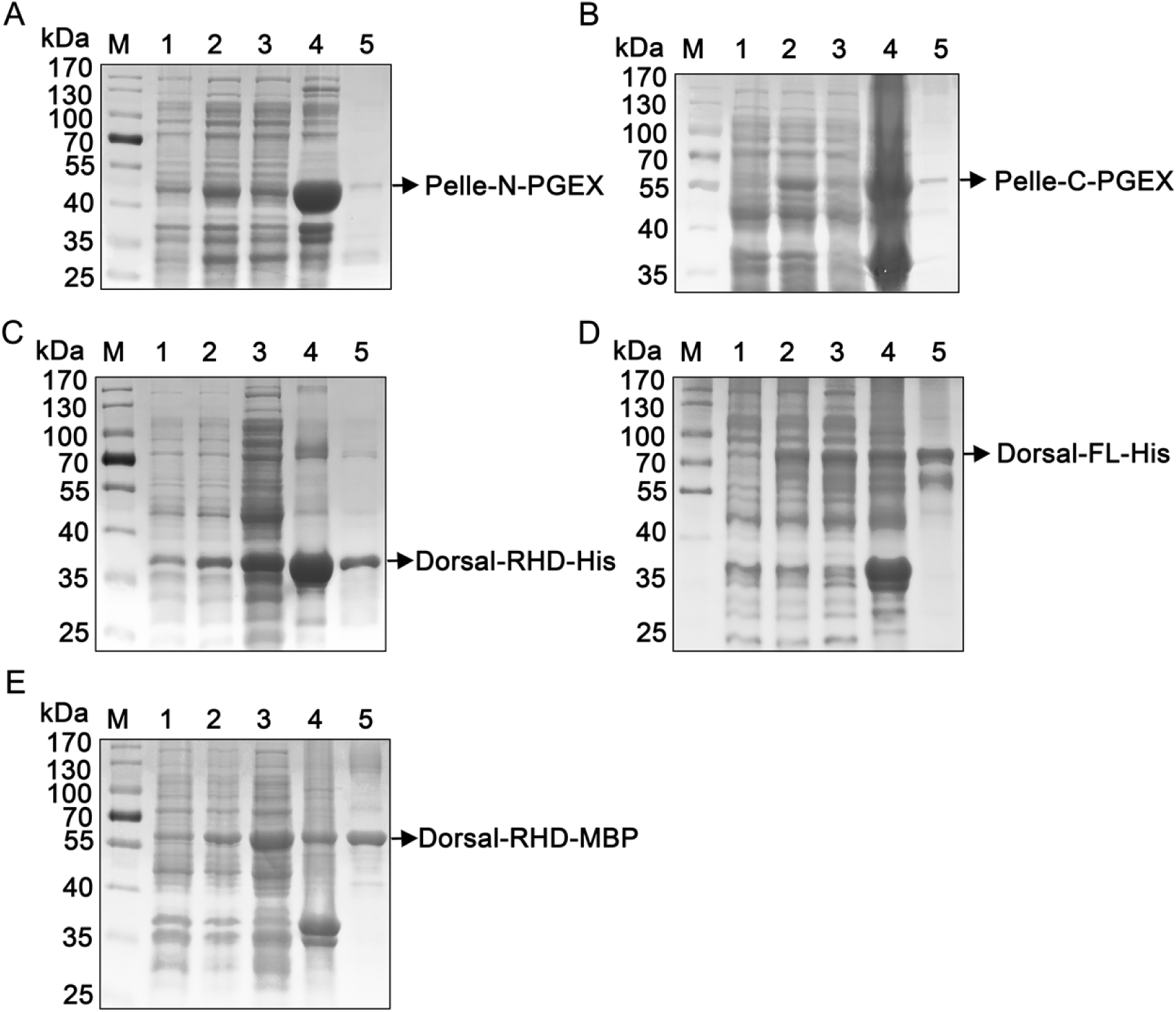
The recombinant protein expression and purification of Pelle-N-PGEX (A), Pelle-C-PGEX (B), Dorsal-RHD-His (C), Dorsal (1-500 aa)-His (D), Dorsal-RHD-MBP (E).

